# A neuromodulatory model for determining the effect of emotion-respiration-cognition coupling on the time-to-respond

**DOI:** 10.1101/2022.03.30.486453

**Authors:** Shogo Yonekura, Julius Cueto, Hoshinori Kanazawa, Noritoshi Atsumi, Satoko Hirabayashi, Masami Iwamoto, Yasuo Kuniyoshi

## Abstract

Respiration and emotional stimuli modulate cognitive ability and the reaction time to generate bodily movement. To understand mechanisms for emotion-respiration-cognition coupling, first, we considered a schematic feed-forward neural network, in which neurons was biased by respiratory-relevant sensory input and the activation function of a neuron was modulated by a neuromodulator, such as norepinephrine (NE). Furthermore, we assumed that the neural model received a stimulus input and generated a response action upon the activity of the output neuron exceeding a certain threshold. Time-to-respond (TTR) was equivalently modulated by the intensity of the input bias and the neuromodulator strength for small action execution threshold; however, it was dominantly modulated by only the neuromodulator for high threshold. Second, we implemented a comprehensive model comprising a cardio-respiration relevant neuromechanical-gas system, a respiratory central pattern generator (CPG), NE dynamics to modulate neurocognitive dynamics, and a locus coeruleus (LC) circuit, which was the primary nucleus for controlling NE. The LC neurons received pCO_2_ or synaptic current from an inspiratory neurons, which resulted in shortened TTR by a stimulus input during inhalation. By contrast, upon receiving pulmonary stretch information, the TTR was shortened by a stimulus input during exhalation. In humans, TTR is shortened when a fear-related stimulus is presented during inhalation, and likewise, TTR is weakly-shortened when surprise-related stimulus is presented during exhalation. Hence, we conclude that emotional stimuli in humans may switch the gating strategies of information and the inflow to LC to change the attention or behavior strategy.

## Introduction

Large portions of cortical and subcortical brain are synchronized with respiratory activities [15, 47], and the synchronization of cardio-respiration and brain activity modulates cognitive activity based on respiration. For example, the accuracy of memory function is significantly modulated by the input timing of stimuli, with respect to the respiratory phase [22, 27, 47]. Furthermore, emotion modulates cognition-respiration coupling; dissimilar emotions differently modulate cognitive functions according to the respiratory phase. The presentation of facial expressions of fear and surprise accelerates the time to respond to the label of an input stimulus during inspiration and expiration, respectively. However, a complete model of emotion-cognition-respiration coupling is not yet available. Nonetheless, researchers have hypothesized two models concerning the mechanism for respiration-cognition coupling [15, 24], namely sensory input(S-)model and neuromodulation(N-) model in Fig. 1. In the S-model, the sensory information of airflow is assumed to be propagated to cortical neurons in several regions via the olfactory bulb [15], therefore, numerous cortical neurons get entrained to the respiratory rhythm. The entrainment of cortical rhythm to respiration is presumably a key mechanism for respiration-cognition coupling. In the N-model, the locus coeruleus (LC), a neural circuit responsible for controlling norepinephrine (NE), projects to several brain regions and gets entrained by respiration; therefore, respiration influences cortical areas via neuromodulators. This article is organized as follows: First, we report an analysis of the behavior of schematic S- and N-models. We introduced a schematic neural model that processes an input signal and executes an action, such as pressing a button during a cognitive decision task for a neuron output crossing a threshold Θ. We demonstrated that sensory input and neuromodulators equally influenced the time-to-respond (TTR) for a stimulus input, with a low Θ value; however, neuromodulators are primarily controller of the TTR, in a high Θ situation. Second, based on the schematic-analytical result, we focused on the N-model in a high Θ condition. We implemented a comprehensive respiration-cognition system by integrating a cognitive decision system, LC-controlled NE, a neurophysical respiration system consisting of a mechanical lung, a O_2_/CO_2_ gas exchange system, and a neural respiration controller [4, 25, 34, 40, 41]. Our model could reproduce the characteristics of TTR modulation according to the respiration phase and different emotional stimuli. We discuss respiratory-relevant inputs to the LC, with gated *p*CO_2_ (the partial pressure of carbon dioxide in blood), pulmonary stretch information (PSR), and inspiratory neural activity, based on differences in the emotional context of the input stimulus.

**Figure 1.**
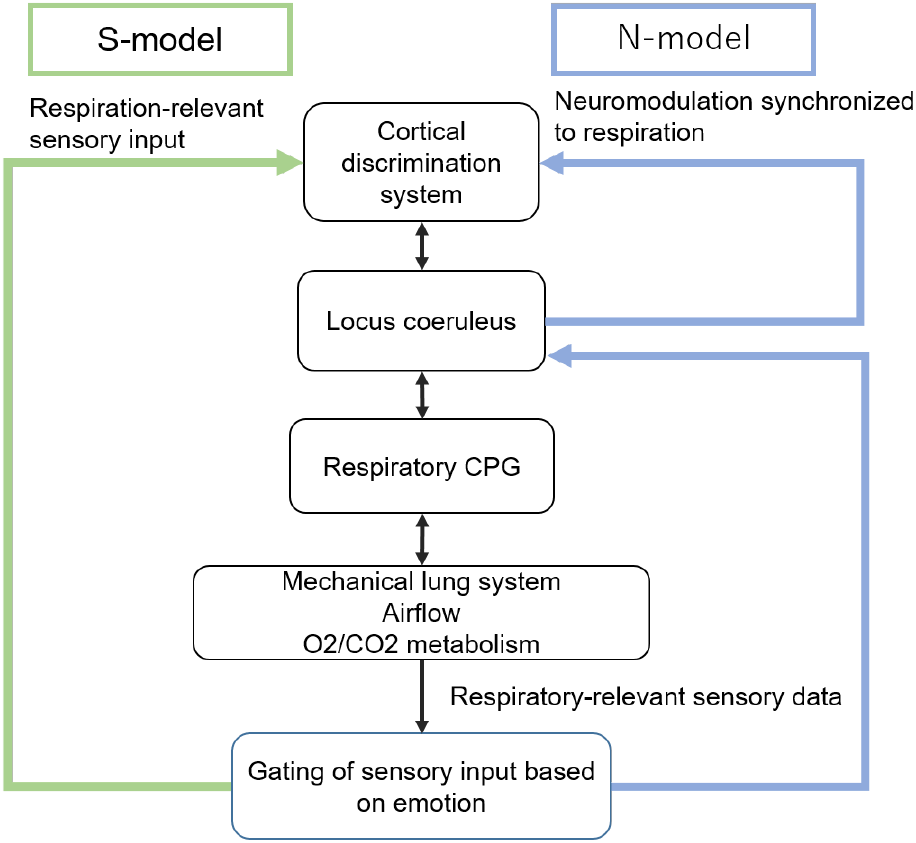
Emotion-respiration-cognition coupling model based on sensory input- (S-) and neuromodulator-based (N-) models. Literature on neurophysiology have proposed two models to account for respiration-cognition coupling, despite limited evidence for the influence of emotion on respiration-cognition coupling. In the S-model, inspiratory information, such as airflow, gets propagated via the olfactory bulb to several cortical regions, therefore, entraining the activity of wide regions of the cortex to the respiratory rhythm. In the N-model, the limbic system is coupled with the respiratory neural system, and controls cortical neurons via neuromodulators. Emotional input may change the gating strategies of the input information to the locus coeruleus. For more details, please refer to text.

## Results

### Schematic analysis

#### Different contributions of sensory input and neuromodulation to TTR

First, we analyzed the behavior of S-/N-models in terms of TTR, (the time-to-execute an action in response to the given stimuli) 2. Herein, we analyzed the contribution of the stationary sensory input *I_sig_* and neuron activation gain *g* to TTR; TTR was computed as the elapsed time for the neuron output responsible for generating an action beyond the threshold Θ, following the stimulus input. The time scale of sensory signals induced by respiration was significantly slower than neuron dynamics, i.e., it was negligible. Therefore, we did not assume fluctuating dynamics induced by respiration for the sake of simplicity. Please refer to Materials and Methods for more details. Furthermore, we assumed that the effect of *I_sig_* and *g* on TTR was correlated to the effects of S- and N-models on TTR, respectively.

The contribution ratio of the S- and N-models to TTR modulation was substantially based on the threshold for an action execution Θ. Fig. 2 depicts the TTR versus *I_sig_* and *g* with several Θ values. For low threshold Θ < 0.5, changes in *I_sig_* and *g* could exert comparably equivalent effects on *TTR*. However, for high threshold Θ, *g* dominantly influenced *TTR*, and *I_sig_* exerted a limited and small effect. For Θ = 0.975, only the activation gain *g* determined the action execution. In other words, the S- and N-models could equally contribute to TTR modulation for low action threshold; however, only the N-model can modulate the TTR for high action threshold.

**Figure 2.**
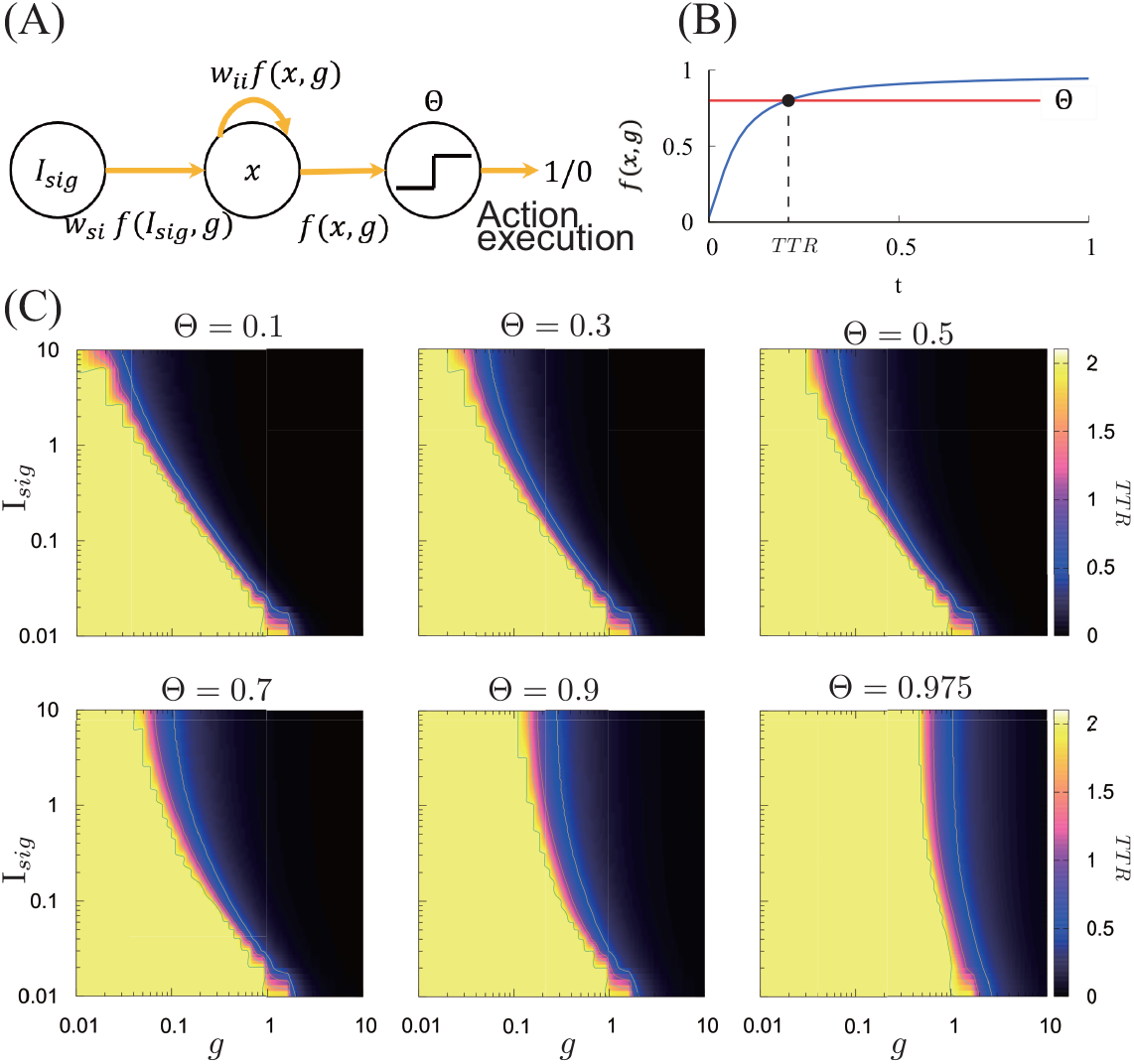
A schematic model to analyze different effects of S- and N-models on the TTR. We considered the effects of the S- and N-models on the TTR by changing *I_sig_* and *g*, respectively. Particularly, we considered a feed-forward neural networks as follows: (A) the first neuron did not demonstrate time-dependent dynamics and generated an output *f*(*I_sig_, g*). The second neuron received the synaptic input *w_si_f*(*I_sig_, g*) and a self-excitable synaptic input *w_ii_f*(*x, g*). We observed the TTR, the time required for *f*(*x, g*) to pass over Θ. (B) the TTR with respect to *I_sig_* and *g* with several Θ. Low action thresholds, such as Θ ≤ 0.5, *I_sig_* and *g*, could equally influence the TTR. However, for high thresholds, such as Θ ≥ 0.9, changes in *I_sig_* induced limited TTR modulation; nonetheless *g* could effectively modify the TTR. We used *w_ii_* = 5, *w_si_* = 5, *D* = 0.1. (Note: time, *x*, and *g* were dimensionless; therefore, did not have units in this schematic model).

The spinal projection of LC modules determines the threshold for a certain kind of emotional action, such as mechanical withdrawal action, in response to pain-inducing stimulus [7, 16]; a high firing rate of the corresponding LC modules increases the threshold for withdrawal action, thereby decreasing the influence of pain-inducing stimulus on the withdrawal action. Therefore, the NE, and not the external sensory input, determines the action generation in a high Θ state, which corresponds to an alert mental state.

### A comprehensive model for respiration-cognition coupling

To conduct more biologically plausible tests for emotion-respiration-cognition coupling, we implemented a multifaceted biological system by integrating a neural CPG for respiration [25], a respiration-relevant lung and gas exchange system [25, 29], a LC neural circuit [2, 33, 43], and a cognitive decision system [43] as shown in Fig. 3-(A, B).

**Figure 3.**
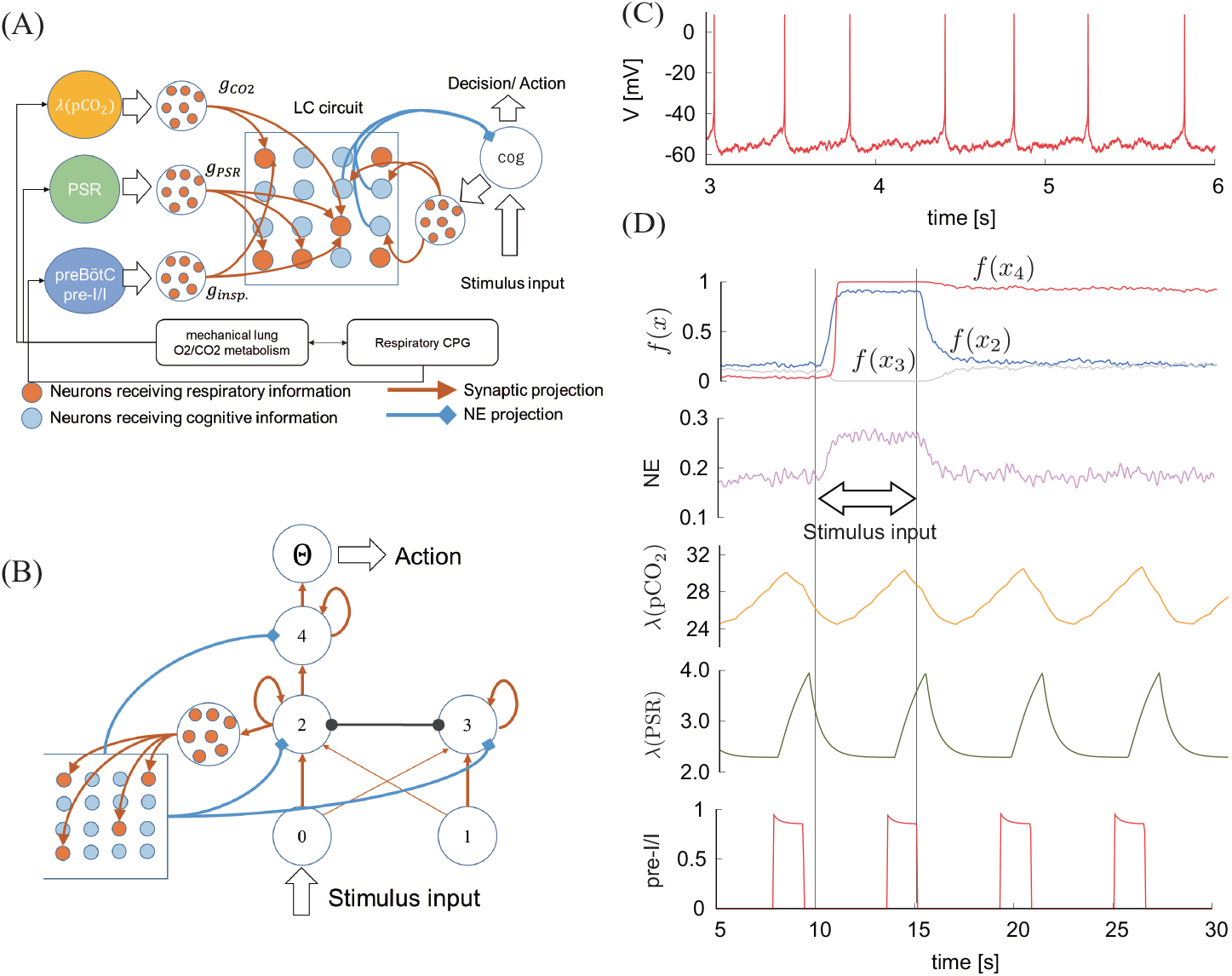
A comprehensive model of respiration system, neuromodulation, and cognition. (A) A schematic model to integrate respiration central pattern generator (CPG), a respiration-relevant lung and gas exchange system, locus coeruleus (LC), and a cognitive decision system; (B) A model of the cognitive neural network and LC. We used a rate-coded neuron model with a sigmoidal activation function for a cognitive decision network and an HH-style spiking neuron model that exhibited slow pacemaker activity for LC, as depicted in panel (C). (C) Slow pacemaker firing of LC neurons. We adjusted the parameters such that LC neuron exhibited 2-3 Hz firing activity. (D) From the top to bottom, the output of cognitive neurons, the amount of norepinephrine (NE) projected to cognitive neurons, low-pass filtered *p*CO_2_ and PSR (i.e., *λ*(*p/rmCO*_2_) and *λ*(PSR)), and the activity of pre-I/I neurons. In this panel, the input stimulus *I_s_* = 1.5 was added to the neuron 0 during *t* = [10, 15]. For more details, please refer to the text.

In our model, the LC presumably received *λ*(PSR) and *λ*(pCO_2_) (low-pass filtered PSR and pCO_2_, respectively), and the synaptic currents from a group of neurons inducing the respiratory inspiration and emotion adjusted the gating of the aforementioned information; emotion supposedly adjusted the weights *g*_CO_2__, *g_PSR_*, and *g_insp_* as shown in Fig. 3-(A). Spatially distributed neurons in the LC form an ensemble, and different ensembles correspond to diverse target cortical areas [28, 42]. Thus, we assumed that randomly selected 100*p* % LC neurons received respiration-relevant information and other (1 − *p*) LC neurons received cognition-related information as shown in Fig. 3-(A, B).

We adjusted the parameters of the LC neuron model to generate spontaneous spikes of approximately 2 Hz as shown in Fig. 3-(C) based on the report that LC neuron generates spontaneous spikes at 0.5-5 Hz [3, 32]. Likewise, we adjusted several system parameters to reproduce human respiration frequency at approximately 0.2 Hz as shown in Fig. 3-(D).

### Different modality of input LC for varied TTR modulation

The TTR is modulated by both emotion and respiration phase [47]. Similar results could be obtained by applying adequate gating to the input to LC neurons. Particularly, the TTR gets reduced by inspiration if the LC coupled with *p*CO_2_ or pre-I/I neuron activity, i.e., *g*_CO_2__ > 0 or *g_insp._* > 0 with *g_PSR_* ~ 0. By contrast, the TTR gets reduced by expiration if the LC is coupled with pulmonary stretch information, i.e., *g_PSR_* > 0 with *g*_CO_2__ ~ 0 and *g_insp._* ~ 0 as shown in Fig. 4. The aforementioned tendency of the TTR behavior, with respect to the respiration phase, did not change with *p*. Moreover, only the magnitude of TTR-modulation by the respiratory phase decreased with *p*. *p* = 0.25 is sufficient to achieve a significant TTR modulation by respiration. Note that these results do not change significantly if we assume LC receives *p*CO_2_ and PSR instead of *λ*(*p*CO_2_) and *λ*(PSR), respectively (see Supplementary Fig. 1).

**Figure 4.**
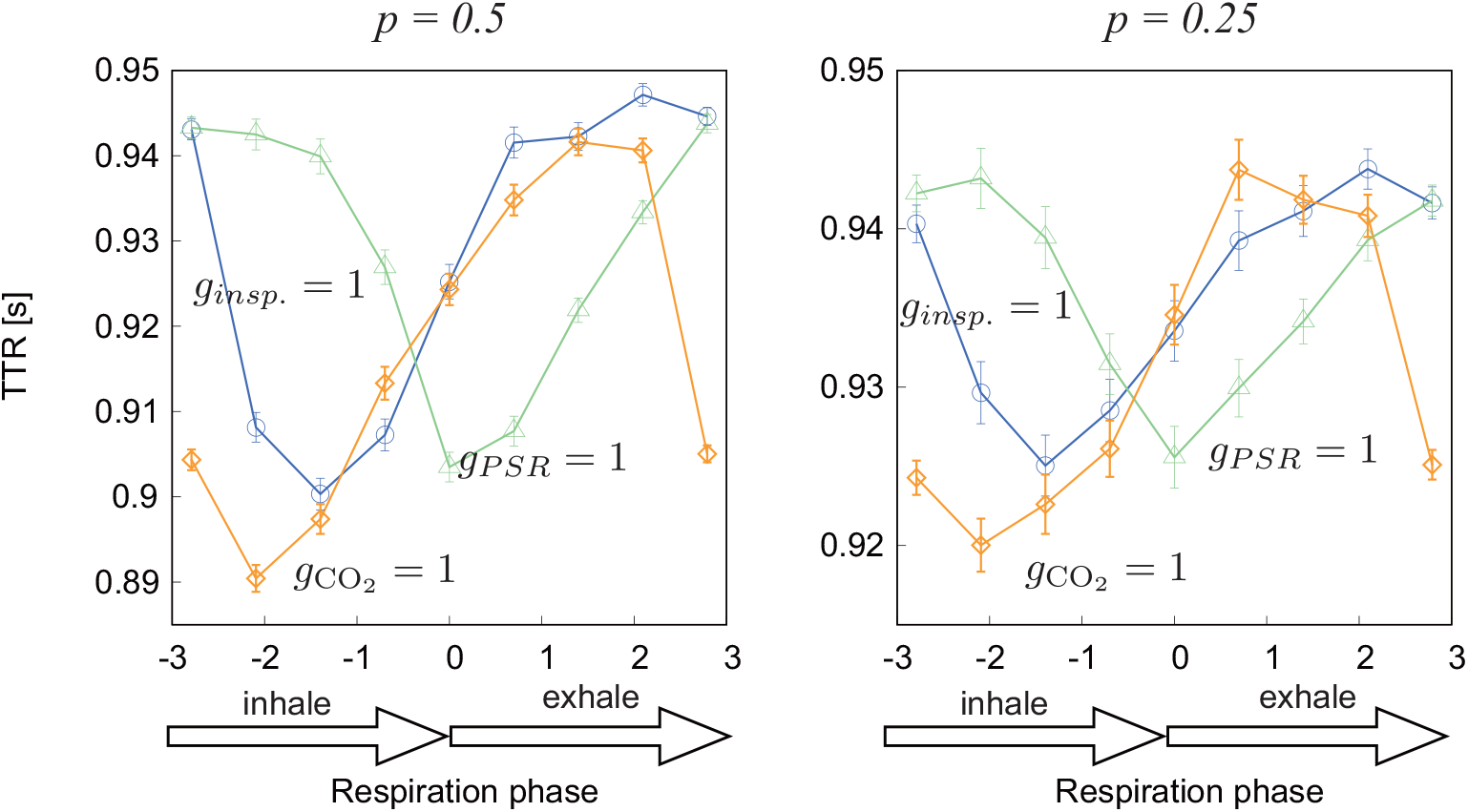
The behavior of TTR modulation by respiration changes by the gating coefficient of respiratory-relevant information. We observed the TTR with weighting variables, namely (*g_insp._*, *g*_CO_2__, *g_PSR_*) = (1, 0, 0), (0, 1, 0), and (0, 0, 1) to gate respiration-relevant information. The labels within the figure are abbreviated, and the description *g_insp._* = 1 denotes (*g_insp._*, *g*_CO_2__, *g_PSR_*) = (1, 0, 0), and similar for others. LC coupled with pre-I/I activity or CO_2_ reduced the TTR by inspiration, For LC coupled with *PSR*, the minimum TTR was generated at the end of inspiration, and the TTR in the expiratory phase was shorter than that in the inspiratory phase. We used Θ = 0.95. Note that inhale corresponds to the respiration phase [−*π*, 0] and exhale corresponds to [0, *π*]. The symbols and bars appearing in the panels denote means and standard errors, respectively.

## Materials and methods

### Schematic model for the TTR modulation

We analyzed TTR modulation by respiration based on the following rate-coded model neuron, with an Arrhenius-type activation function, as follows:

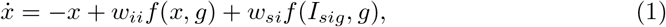

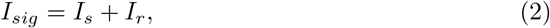

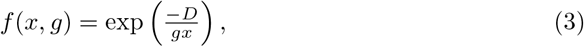

where *I_s_* denotes the input (such as the image of a face expressing fear or surprise), to which an agent must generate an answering action (e.g., decide if the presented image is categorized as fear or surprise); and *I_r_* denotes respiration-relevant information. The timescales of respiratory-relevant sensory information and neuromodulation were significantly slower than the cortical neuron dynamics. Therefore, we could apply an adiabatic limit. Thus, we assumed that *I_s_* and *I_r_* were static and independent of time. The coefficients *w_ii_* and *w_si_* denote weights for a self-activation connection and sensory input, respectively. Furthermore, *f*(*x, g*) exhibited a sigmoidal curve in response to *x*, and *g* determined the non-linearity of the function. *f*(*x, g*) approached a step function and exhibited linear behavior with a large *g* and a small *g*, respectively. Likewise, *D* denotes a scaling factor; A small *D* displayed high non-linearity, whereas a large *D* displayed small non-linearity. The effects induced by changes in *I_sig_* and *g* corresponded to those of the S- and N-models on the respiration-cognition coupling, respectively.

### An overview of the integrated respiration-LC-cognitive system

To conduct a biologically plausible experiment in the paradigm of TTR modulation by emotion and respiration phase, we integrated the previously proposed respiration system, neuromodulation system, and cognitive system [2, 25, 29, 43]. Our comprehensive system consisted of a respiratory central pattern generator (CPG) and a O_2_/CO_2_ gas exchange system [25], a lung mechanical system with a non-linear muscle dynamic [29], a LC circuit, and a NE-modulated cognitive decision system [2, 43], as shown in Fig. 3-(A).

LC neurons received respiratory information relevant to pCO_2_ and synaptic signals from the pulmonary stretch receptor (PSR) and the pre-Bötzinger inspiratory pre-I/I neurons. Cardiorespiratory information, such as pCO_2_ or pulmonary stretch information is monitored by the nucleus tractus solitari (NTS), which sends direct and indirect synaptic connections to the LC and amygdala [8, 21, 45]. Furthermore, the amygdala and LC are mutually connected by several types of neurotransmitters, such as noradrenergic synapses, GABAergic, and corticotropin-releasing factor [5, 23, 44, 48]. In our model, the respiration-relevant information *λ*(pCO_2_) ( low-pass filtered pCO_2_), *λ*(PSR) (low-pass filtered PSR), and pre-I/I neuron activity were presumably bypassed by the ensembles consisting of five Hodgkin-Huxley (HH) neurons, and were sent to the LC neurons via synaptic connections, with gating weights *g*_CO_2__, *g_PSR_*, and *g_rp_*. Therefore, LC received the cardio-respiratory related information in the following form: *g_PSR_I_PSR_* + *g*_CO_2__*I*_CO_2__ + *g_rp_I_rp_* + *g_cog_I_cog_*. Moreover, the NTS-amygdala-LC circuit should provide a gating function for the variables, namely *g_PSR_*, *g*_CO_2__, and *g_rp._ λ*(pCO_2_) denotes the application of low-pass filter to pCO_2_ as *τ_λ_ dλ*(pCO_2_)/*dt* = − *λ*(pCO_2_) + pCO_2_ with *τ_λ_* = 1000 [ms], and the same low-pass filter is applied to PSR.

However, there is no concrete model of the connection topology among sensory neurons, LC neurons, and NE target neurons [32]. Therefore, we assumed that randomly selected 100*p* % of neurons in the LC received respiration-relevant information, based on recent reports that spatially distributed neurons in the LC form an ensemble and different ensembles modulate varied cortical areas [28, 32, 42]. We further assumed that other (1 − *p*) neurons in the LC received the activity of a specific cognitive neuron 2 as shown in Fig. 3-(A, B), and projected NE to modulate the gain *g* of the activation function. The LC neuron was considered responsible for modulating NE of the cognitive system, and did not receive respiration information directly.

A cognitive decision system consists of four artificial neurons (1 − 4) as shown in Fig. 3-(B). Neurons 0 and 1 send input- and distractor-signals to decision neurons 2 and 3, respectively. Neurons 2 and 3 are mutually inhibiting, and the output of neuron 2 is passed to the ensemble consisting of 20 HH neurons. Furthermore, the ensemble sends synaptic current to (1 − *p*) of the LC neurons. The decision action was supposedly generated when the output of neuron 4 exceeded the threshold Θ. Only neuron 2 sends information to the LC, but all neurons 2, 3, and 4 are affected by NE.

As shown in Fig. 3-(C), the parameters of LC neuron are adjusted such that the neuron generates slow pacemaker firing at 2 ~ 3 Hz. Furthermore, the panels depicted in Fig. 3-(D) denote (from top to bottom), the output *f*(*x, g*) of cognitive neurons, the amount of NE projected to cognitive neurons, *λ*(pCO_2_), *λ*(PSR), and the activity of pre-I/I neuron. The phase of respiration *ϕ* is defined as [−*π, π*], such that the beginning of inspiration and end of expiration were −*π* and *π*, respectively. Note that *p*_CO_2__ peaked at the beginning of inspiration at *ϕ* = −*π*, and *PSR* peaked at the end of inspiration at *ϕ* = 0.

#### A respiratory neural system, gas exchange system, and lung mechanical system in a human scale

Our respiration system was based on a previously reported model [25], which consisted of a respiratory CPG, O_2_/CO_2_ gas exchange system, and lung mechanical system. However, we adjusted the parameters of the model [25, 26] to reproduce the respiration of an adult male. First, we modified the mechanical lung system from a linear mass-spring model to a non-linear mass-spring model [29]. Furthermore, we made the following changes: *E_lung_* = 3.6595 [mmHg/L], *V_c_* = 0.07 [L], *D*_*co*2_ = 7.08 × 10^−6^ [L/mmHg/ms], and *D*_*o*^2^_ = 3.5 × 10^−7^ [L/mmHg/ms]. The cardiovascular parameters [25] were modified as *H_T_* = 60/72 [s], and the airflow between the mouth and alveoli was computed as 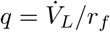, with a flow resistance *r_f_* = 1000[*mmHg* · *ms/L*].

#### A model for respiration-modulated LC circuit

An LC neuron exhibits slow pacemaker firing [2, 43]. Despite propositions for a detailed computational model for a LC neuron with a 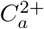 ion channel and *p*CO_2_/*H*^+^ chemoreceptor [33], we used a simplified model that could generate slow pacemaker firing. Our model was only based on the persistent Na^+^ current *I_NaP_* and slowly activated *K*^+^ current *I_KS_* [18]. The neuron model was described as follows:

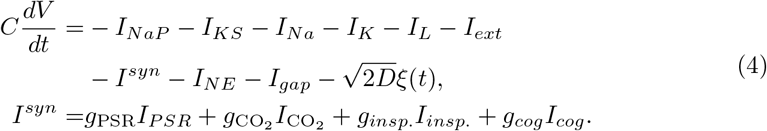

The behavior of the ionic channel has been described in the style of the HH neuron model [18]

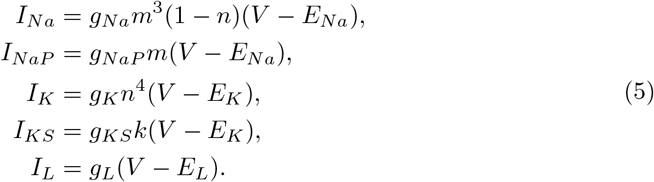

The conductance for each ion channel is described as follows:

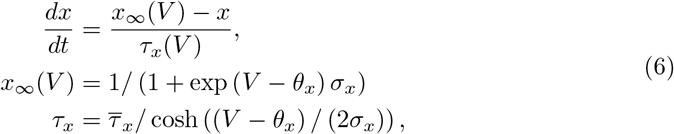

where *x* denotes either *m, n*, or *k*. We used *E_Na_* = 50 [mV], *E_K_* = −85 [mV], *E_L_* = −59 [mV], *g_K_* = 11.2 [nS], *g_KS_* = 5.6 [nS], *g_Na_* = 28 [nS], *g_NaP_* = 2.5 [nS], *τ_KS_* = 600 [ms], *θ_n_* = −29 [mV], *σ_n_* = −4 [mV], 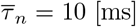, *θ_m_* = −40 [mV], *σ_m_* = −6 [mV], *θ_k_* = 38 [mV], and *θ_m_* = 40 [mV].

An LC neuron has CO_2_ chemoreceptor [1, 11, 12, 33]; however, neuron dynamics are significantly modulated by CO_2_, particularly in hypercapnia. We ignored the chemoreceptor for *p*CO_2_ in LC neurons, and considered emotion-respiration-cognition coupling only in low CO_2_ condition.

Neighboring eight neurons in the LC were electrically connected by the gap-junction and mutually inhibited by local NE diffusion [2, 43]. Local inhibitory current by NE was computed as 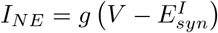 with 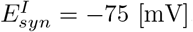. *g* is the effective conductance of locally diffused NE (see the Eq. 7 described later). The membrane current induced by the gap junction was computed as *I_gap_* = ∑_*j*_ *g_gap_*(*V_i_* − *V_j_*) with *g_gap_* = 0.04 [nS].

#### Modulation of cortical neurons by LC-innervated NE

LC neurons diffuse NE to the cognitive neurons and modulate the activation gain *g* [2, 7, 13, 32, 37, 43]. The amount of NE diffused to the cognitive system and the modulatory dynamics for the gain *g* are described as follows: using the second order delayed system [13],

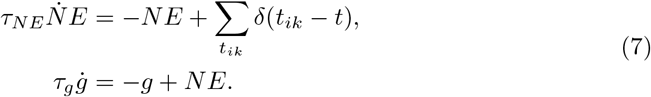

where *δ*(*t_ik_*) indicates the time *t* when the ith neuron in the LC emitted a spike. We used *τ_NE_* = 100 [ms] and *τ_g_* = 300 [ms]. Previous researchers [13] used *τ_NE_* = 300 [ms] and *τ_g_* = 100 [ms]; however, (*τ_NE_, τ_g_*) = (300, 100) and (100, 300) exhibited similar behavior, despite different linear scales.

#### A neuro-ensemble model to transform respiratory-relevant information and cognitive activity into synaptic currents to LC neurons

LC neurons receive excitatory synaptic connections from respiratory CPGs [46], This effect is described by the term 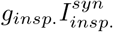, where 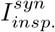 denotes the synaptic current from the pre-Bötzinger inspiratory pre-I/I neuron and *g_insp._* is a gating variable.

Synaptic current *I_z_* to a LC neuron from a neuron ensemble that received either *PSR*, pCO_2_, *insp.*, or *cog.* was computed using the spike-trains *t_k_* as follows:

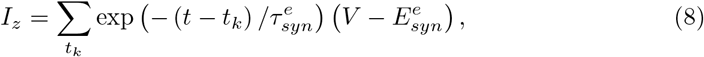

where 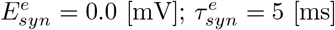; and *z* denotes either *PSR*, CO_2_, *insp*., or *cog*..

We modeled a neuron ensemble receiving *PSR*, CO_2_, or pre-I/I neuron activity using an HH-style neuron as follows:

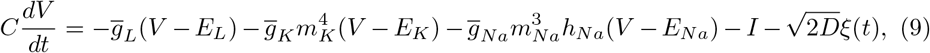

where *C* = 36 [pF],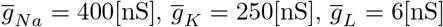, *E_Na_* = 55[mV], *E_K_* = −94[mV], *E_L_* = −60 ± 1.2*σ* [mV] (*E_L_* was based on Gaussian distribution, with mean −60 [mV] and variance 1.2), *ξ*(*t*) is the Gaussian noise with unit intensity, and *D* is the noise intensity. Furthermore, *m_K_*, *m_Na_*, and *h_Na_* denoted ionic channel activation function, and we used *τ_mz_dm_z_*/*dt* = *m_∞z_*(*V*) − *m_z_*, where *z* = *K* or *Na*, and *τ_hz_dh_z_/dt* = *h_∞z_*(*V*) − *h_z_*. The stationary values for these ionic channel behaviors were based on previous reports [4] as follows: *m_∞N_a__* = 1/(1 + exp(−(*V* + 34)/7.8)), *h_∞_N_a_* = 1/(1 + exp((*V* + 55)/7)), *m_∞K_* = *α_∞K_*/(*α_∞K_* + *β_∞K_*), *α_∞K_* = 0.01(*V* + 44)/(1 − exp(−(*V* + 44)/5)), and *β_∞K_* = 0.17 exp(−(*V* + 49)/40). Furthermore, we used the time constants as follows:

*τ_hNa_* = 8.456/ cosh((*V* + 67.5)/12.8) [ms], and *τ_mK_* = 3.5/ cosh((*V* + 40)/40)[ms]. We assumed that *m_Na_* = *m_∞N_a__* (*τ_mNa_* = 0).

A neuron ensemble to convert the signal of PSR, pCO_2_, or pre-I/I neuron activity into the corresponding synaptic current to LC neurons consists of five neurons with *I_b_* = −35 [pA] and *D* = 60 [pA]. In particular, the PSR and pCO_2_ signals are processed by a lowpass filter and converted to bias currents as *I* = 520 − 5.625*λ*(PSR) + *I_b_*, and *I* = 70 − 3*λ*(pCO_2_) + *I_b_*. Note that *λ*(*z*) denotes a lowpass filter is applied to *z*. The pre-I/I signal is passed to a HH neuron ensemble as a synaptic current as 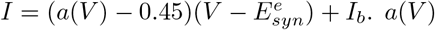 denotes the activation function of a neuron as follows [25]:

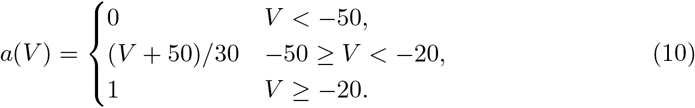

#### A model for cognitive decision-making and action execution

We used a rate-coded neuron model to implement a system for cognitive decisions and action execution [2, 43]. The cognitive system consisted of two input neurons and two decision neurons as shown in Fig. 3-(B). The dynamics of the *i*th neuron was described as follows:

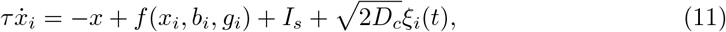

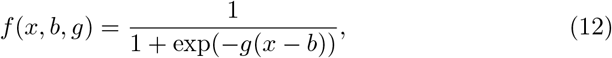

where *f*(*x, b, g*) denotes the sigmoidal function with gain *g* and bias *b*. *g* was modulated by the LC-innervated NE, and we used *b* = −3.0 and *b* = −1.5 for the decisional action neuron *i* = 4 and other cognitive neurons, respectively. *D_c_* denotes the intensity of a Gaussian noise, and we used *D_c_* = 10^−5^.

Each input neuron received an input *I_s_* and sent an excitatory input to the corresponding decision neuron. *f*(*x*_4_, *b, g*) denoted the output of a decision neuron and executed the corresponding motor action upon passing the threshold Θ (for example, in a previous experiment [47], the action involved pressing a button labeled as either fear or surprise). Each decision neuron has a self-excitatory synapse connection, and projects an inhibitory synapse to the opposing decision neuron. *f*(*x, b, g*) approached a threshold-like shape, with respect to *x* by *g* → ∞, and a linear function, with respect to *x* by *g* → 0.

We added *I_s_* = 1.5 to an input neuron *i* = 0 as shown in Fig. 3. *I_s_* presumably corresponded to a visual stimulus, such as facial expression in physiological experiments [47]; however, we assumed that *I_s_* was identical and = 1.5, regardless of the difference in face expression (either fear or surprise). Rather, the difference of emotional valence of a stimulus modified the gating weights for the input to LC neurons.

#### Computing the TTR versus input-timing in terms of the respiratory phase

To obtain the results in Fig. 4, we randomly modified the time *T_s_* to add an input stimulus *I_s_* = 1.5 to the cognitive system, and observed the time *T_r_* when the activity of the action-output neuron *i* = 4 exceeded Θ. Thus, *TTR* is computed as TTR = *T_r_* − *T_s_*. The time of stimulus input *T_s_* was converted to the respiratory phase *ϕ* by using the Hilbert transformation to *λ*(pCO_2_) and shifting by 1/4*π*. This is because 1/4*π* delayed the *ps* signal to the phase of the airflow.

## Discussion

### Cognitive modulation by respiratory-relevant sensory signals and neuromodulation

Our results imply that in the presence of a high threshold-like mechanism to execute a bodily action, the sensory signal input rarely influences the action execution timing. Moreover, the neuromodulator is a dominant information transmitter to generate a bodily action. Previous computational studies have demonstrated that respiratory-relevant sensory input via the olfactory bulb to fewer cortical neurons could induce the overall cortical synchronization to respiration [15]. Our findings emphasized that the sensory input-based synchronization of cortical neurons to respiration may not be sufficient for the timing of bodily action generation modulated by respiration.

### Respiratory signal bridges information transmission in an LC circuit through synchronization

In our LC model, respiratory information was supposedly transmitted to spatially distributed LC neurons. Fig. 4 depicts that greater than 1/4 of the LC distributed neurons received respiratory information. Thus, other LC neurons that did not receive respiratory information were also entrained to respiration. The overall LC entrainment based on the signal input to fewer neurons was based on a similar mechanism to neuropercolation [15, 19], where a limited number of locally synchronized neuron clusters entrain the overall neurons.

### Emotion may control the coupling of different interoceptive dynamics and cognitive functions

In the human respiration-cognition coupling experiment [47], the visual input of different facial expression (e.g., fear or surprise) may have led to the distinct coupling of LC with PSR and pCO_2_ or inspiratory neural activity as shown in Fig. 4. We speculate that emotion may provide an adaptive gating of information in flow to the neuromodulator-controlling nucleus, such as the LC and substantia nigra, to realize an adequate coupling of interoceptive information (e.g., cardiovascular, respiratory, and gastrointestinal information) and cognitive function for a reasonable purpose.

The coupling of interoceptive-exteroceptive inference facilitates the generation of an adaptive behavior to regulate homeostasis [39]. However, this theory may not sufficiently account for the switches in the coupling of cognition and inhalation/exhalation depending on the emotion. We speculate that interoceptive-cognitive coupling should provide adaptability to survival or social tasks for an embodied agent. For example, speeding up the reaction to a stimulus during inhalation in a fearful situation may generate a sharp and large adaptive body motion to realize fight-or-flight behavior, in terms of the metabolism of *O*_2_ in muscles. This warrants understanding the benefits of interoception-cognition coupling, in terms of the survival or social activity of an embodied agent.

### A Mechanism for the adaptive interoception-cognition coupling

Our finding that emotion provides adaptive information gating to realize adequate intero-extero information coupling was consistent with the finding that the amygdala-LC circuit provides gating for early sensory processing [14]. Two famous and traditional principles of emotions are possible candidates for the adaptive gating mechanism, namely, adaptive gating initiated by the central nervous system (CNS) based on the Cannon-Bard theory [6] and a body-initiated peripheral mechanism based on the James-Lange theory [17, 20].

Recent findings that attention adaptively controls the intensity of cardiovascular and cortical electroencephalogram coupling [30, 31] may support the idea of CNS-initiated control of interoception-cognition coupling. Intuitively, the body-initiated peripheral mechanism for the adaptive control of interoception-cognition coupling appears biologically plausible, for example, surprise often enhances quick and large inhalation. The breath-holding [10] bodily reaction accompanying surprise would lead to a sudden and large pulmonary stretch, which would further cause large interoceptive prediction error in the anterior cingulate cortex or LC [36]. Therefore, it would increase the importance weighting of the PSR input to the LC.

Recently, several studies have reported that adequate voluntary breathing is capable of relieving emotional and mental distress, such as post-traumatic stress disorder and hypertension [9, 22, 35, 38], and may provide further evidence supporting the idea of peripheral-initiated control of interoception-cognition coupling. Voluntary deep breathing generates a large PSR and low pCO_2_, and the aforementioned sudden changes in respiratory-relevant information helps evading the coupling of cognition and pCO_2_ or pre-I/I inspiratory information(which corresponded to the context of fear in our experiment), and transiting to the interoception-cognition coupling typical to a comfortable mental state. This necessitates better understanding of the exact mechanism of emotion regulation by adequate voluntary breathing.

## Conclusion

In this study, we considered a mechanism for emotion-cognition-respiration coupling. We conducted an analysis using a schematic neural model in which the response to a stimulus input was generated when the output exceeded a certain threshold. For high threshold, the amount of neuromodulator NE influenced the TTR more effectively than the input stimulus intensity. We conducted comprehensive analyses using a more biologically detailed system consisting of a respiration system, LC, NE, and a cognitive system. The coupling of LC with pCO_2_ or inspiratory neuron activity reduced the TTR upon presenting a stimulus during inhalation. By contrast, the coupling of LC with PSR reduced the TTR upon presenting a stimulus during exhalation. Our results were consistent with psychological experiments using human participants, which demonstrated that the TTR gets reduced on presenting a fearful face as a visual stimulus during inhalation. Our results imply that distinct emotions may provide different gating strategies for the information input to the LC, and may generate dissimilar interoception-cognition coupling to optimize survival or social ability.

## Acknowledgments

We would like to express our gratitude to Dr. Yoshiyuki Ohmura (the University of Tokyo), Dr. Hiroyuki Sakai (Toyota Central R&D Labs., Inc.),and Mr. Daisuke Yamada (Toyota Central R&D Labs., Inc.) for their valuable discussions.

