## Supplementary figures and images for "A neuromodulatory model for determining the effect of emotion-respiration-cognition coupling on the time-to-respond"

### Supplementary Figure 1

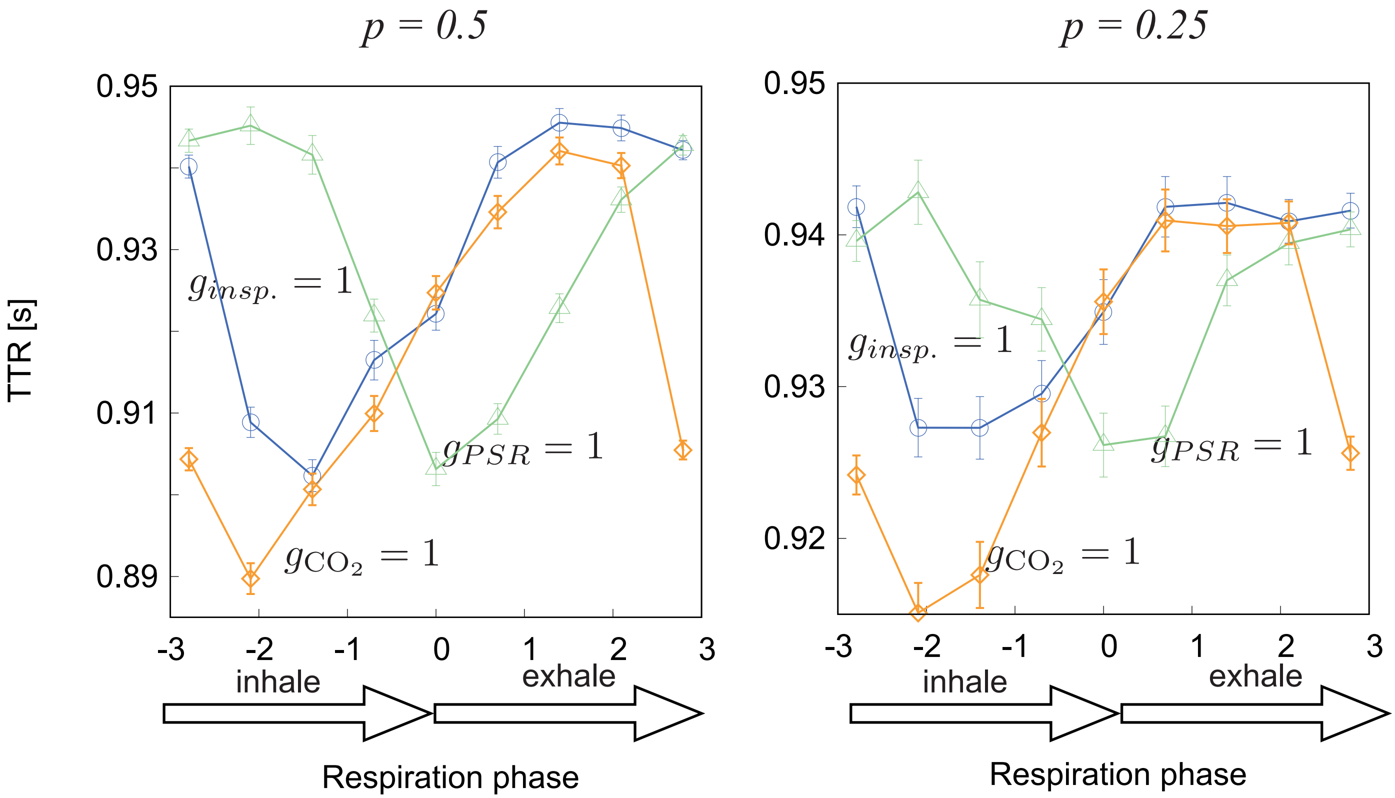
